# Progerin Can Induce DNA Damage in the Absence of Global Changes in Replication or Cell Proliferation

**DOI:** 10.1101/2024.07.02.601729

**Authors:** Liza A. Joudeh, P. Logan Schuck, Nina M. Van, Alannah J. DiCintio, Jason A. Stewart, Alan S. Waldman

**Affiliations:** Department of Biological Sciences, University of South Carolina, Columbia, SC 20208; Department of Biology, Western Kentucky University, Bowling Green, KY 42101

**Keywords:** Hutchinson-Gilford Progeria Syndrome, Progerin, Aging, DNA damage, DNA double-strand break repair, Genomic instability, Nuclear lamina

## Abstract

Hutchinson-Gilford Progeria Syndrome (HGPS) is a rare genetic condition characterized by features of accelerated aging, and individuals with HGPS seldom live beyond their mid-teens. The syndrome is commonly caused by a point mutation in the LMNA gene which codes for lamin A and its splice variant lamin C, components of the nuclear lamina. The mutation causing HGPS leads to production of a truncated, farnesylated form of lamin A referred to as “progerin.” Progerin is also expressed at low levels in healthy individuals and appears to play a role in normal aging. HGPS is associated with an accumulation of genomic DNA double-strand breaks (DSBs) and alterations in the nature of DSB repair. The source of DSBs in HGPS is often attributed to stalling and subsequent collapse of replication forks in conjunction with faulty recruitment of repair factors to damage sites. In this work, we used a model system involving immortalized human cell lines to investigate progerin-induced genomic damage. Using an immunofluorescence approach to visualize phosphorylated histone H2AX foci which mark sites of genomic damage, we report that cells engineered to express progerin displayed a significant elevation of endogenous damage in the absence of any change in the cell cycle profile or doubling time of cells. Genomic damage was enhanced and persistent in progerin-expressing cells treated with hydroxyurea. Overexpression of wild-type lamin A did not elicit the outcomes associated with progerin expression. Our results show that DNA damage caused by progerin can occur independently from global changes in replication or cell proliferation.

## Introduction

Genomic stability in mammalian cells relies on a cell’s ability to successfully correct a multitude of forms of DNA damage that arise each day. One type of damage that cells must contend with is a DNA double-strand break (DSB). A DSB can be generated by chemical or radiological insult, form spontaneously from other DNA lesions, or form at stalled or collapsed replication forks. It is important that DSBs be repaired efficiently and accurately to avoid potentially deleterious chromosomal rearrangements or mutations.

Mammalian cells have two general types of DSB repair pathways at their disposal: homologous recombination (HR) and nonhomologous end-joining (NHEJ) [reviewed in 1-10]. The salient difference between these two broadly defined repair strategies is that HR utilizes a template sequence to maintain or restore genetic information to the DSB site that may otherwise be lost through strand degradation, whereas NHEJ involves no template in the rejoining of DNA ends. HR is thus generally considered to be accurate while the NHEJ pathway, despite its potential for healing DSBs accurately, is comparatively error-prone and not infrequently produces deletions or insertions.

The efficient and accurate execution of DNA repair pathways is of paramount importance in maintaining genomic stability. However, DNA repair pathways can become corrupted and, in turn, generate genomic instability which can take a variety of forms. The association between cancer and aberrant DSB repair is well-documented in the literature [11–13]. Genomic instability has also been associated with the process of aging. Increased levels of damage, mutation, and large-scale chromosomal abnormalities such as translocations, insertions and deletions have been observed with increasing age in humans, and other organisms [14–27]. There is much evidence in the literature that the origin of the increase in genomic instability that accompanies aging is a decrease in the effectiveness of a variety of DNA repair pathways. Alterations in NHEJ were reported in rat brain during aging [20], and studies with mice have suggested that the fidelity of DSB repair diminishes with age [21]. Both the efficiency and fidelity of DNA end-joining has been observed to decrease as human fibroblasts approach senescence [22]. Chromosomal DSBs accumulate in human cells approaching senescence, and it has been suggested that DSBs may be involved directly in the induction of senescence [14]. As the integrity of the genome is progressively compromised over time, general cellular functions would be expected to be disrupted. In addition, cell number would gradually be reduced as cells are lost due to apoptotic responses to unrepaired DNA lesions. Loss of cell number and associated tissue depletion would further contribute to loss of biological functions. Thus, the accumulation of mutation and DNA damage is viewed as a possible basis for, or at least a significant contributor to, the aging process.

As might be expected, genetic disorders that produce clinical features of premature aging (progeria) are often associated with DNA repair defects and associated genomic instability [15–19, 23–27]. Hutchinson–Gilford Progeria Syndrome (HGPS) is one such genetic syndrome that leads to accelerated aging [reviewed in 28]. Individuals with HGPS rarely live beyond their teens. HGPS is most commonly caused by a point mutation in the LMNA gene which normally codes for lamin A and its splice variant lamin C. The LMNA mutation associated with HGPS leads to increased usage of a cryptic splice site which leads to the production of a truncated form of lamin A referred to as “progerin.” Significantly, it has been learned that progerin is in fact expressed at low levels in healthy individuals and appears to play a role in the normal aging process [29–32]. Unlike wild-type (wt) fully processed lamin A, progerin retains a farnesyl group and a methyl group at its carboxy terminus. These modifications cause progerin to largely remain associated with the inner nuclear membrane rather than localize to the nuclear lamina where lamin A normally resides.

In HGPS, progerin overexpression has a severe effect on the nuclear lamina which, in turn, has severe effects on nuclear architecture and function. The nuclei of HGPS cells are characteristically misshapen and display blebs and invaginations. The altered nuclear structure imparts important changes to numerous nuclear functions and profoundly alters chromatin organization. One impact of progerin expression in HGPS cells is an accumulation of DSBs and increased sensitivity to DNA damaging agents [33–40]. Studies have revealed that recruitment of repair proteins, most specifically those involved in HR repair (including Rad 50, Rad51, NBS1, and MRE11), to the site of a DSB is delayed in HGPS [36,37]. In accord with such findings, several studies have provided evidence of an enhancement of NHEJ with a concomitant suppression of HR in association with progerin expression [41–44]. We have developed an experimental system using cultured mammalian cells containing an integrated DSB repair reporter substrate into which we can induce a DSB by expression of endonuclease I-SceI. Using this model system, we directly demonstrated [45] that repair of a genomic DSB is indeed shifted away from HR and toward NHEJ in cells expressing progerin. Additionally, repair by HR is shifted away from crossovers and toward gene conversions in cells expressing progerin [45]. We recently also reported [46] that the precision of DNA end-joining repair of a genomic DSB is reduced in the presence of progerin expression. A unifying theme of our findings is that progerin brings about an apparent diminution of a cell’s ability to anneal complementary terminal DNA sequences at the site of a DSB, and this dysfunction is likely to reduce the efficiency of DSB repair and destabilize the genome.

Despite an expanding knowledge of the impact that progerin expression has on the nature of DSB repair events, many unknowns remain, including the lack of a firm understanding of the temporal sequence of events that may be responsible for the genesis of the DSBs that accumulate in the genomes of HGPS patients. There is evidence that much progerin-induced damage occurs during S-phase of proliferating cells [reviewed in 47]. Among its various roles, lamin A normally helps recruit proliferating cell nuclear antigen (PCNA), DNA polymerase delta and other factors to replication forks. Progerin expression in HGPS interferes with this recruitment, often leading to replication fork stalling and collapse [33–36, 47–51]. Such collective observations have led to the paradigm that stalled and subsequently collapsed replication forks are a direct and primary source of endogenous DNA damage in HGPS. Matters are made worse by the mislocalization and binding of XPA to sites of breakage at collapsed replication forks [34,35,48]. Mislocalization of XPA is believed to suppress DSB repair by stearic hindrance of the appropriate repair proteins. The essence of these collective observations is that problems with replication fork progression set in motion a series of events that results in genome damage and ultimately cellular demise.

In the current work, we used our model system to investigate the levels of endogenous damage in cells expressing progerin compared to levels of damage in cells not expressing progerin. We now report that, consistent with other studies, progerin expression was associated with significantly increased endogenous chromosome damage but, notably, we observed progerin-associated genomic damage in an immortalized cell line that exhibited neither an increase in cell doubling time nor an alteration in cell cycle profile compared with control cells. Additional experiments suggested that progerin-expressing cells exhibit a reduced ability to recover from replication fork stalling and/or a particular sensitivity to alteration of nucleotide pools. Our model system, in a straightforward way, challenges a paradigm in which obstructed replication necessarily serves as the initiating source of progerin-inflicted endogenous DNA damage.

## Materials and Methods

### General cell culture

All cell lines were derived from normal human fibroblast cell line GM637 which was obtained from the NIGMS. The GM637 cell line is immortalized by SV40. Cells were cultured in alpha-modified minimum essential medium supplemented with 10% fetal bovine serum. All cells were maintained at 37°C in a humidified atmosphere of 5% CO_2_.

### DNA recombination and repair substrate

Plasmid pLB4 is a recombination and DSB repair reporter substrate and was described previously [52,53]. Briefly, pLB4 contains a gene comprised of herpes simplex virus type 1 thymidine kinase (tk) sequence fused to a neo gene sequence. The tk-neo fusion gene is disrupted by a 22 bp oligonucleotide containing the 18 bp recognition sequence for endonuclease I-SceI. The substrate pLB4 also contains a “donor” tk sequence that shares about 1.7 kb of homology with the tk portion of the tk-neo fusion gene.

### Cell lines

As previously described [54], substrate pLB4 was stably integrated into the genome of human cell line GM637 cells to produce cell line pLB4/11 containing a single integrated copy of pLB4. To produce a derivative of pLB4/11 that constitutively expresses GFP-progerin, cell line pLB4/11 was stably transfected with plasmid pBABE-puro-GFP-progerin (Addgene plasmid # 17663) as described [45]. This latter plasmid, a gift of Tom Misteli from the National Cancer Institute, expresses progerin as a GFP fusion protein under the control of a Moloney murine leukemia virus LTR promoter. The derivative of cell line pLB4/11 that expresses GFP-progerin is named pLB4-progerin. A derivative of pLB4/11, named pLB4-GFP, that constitutively expresses GFP was also previously produced as described [45].

To produce a derivative of pLB4/11 that constitutively expresses GFP-wt lamin A, plasmid pBABE-puro-GFP-wt Lamin A (Addgene plasmid #17662, gift of Tom Misteli) was transfected into pLB4/11 cells in the same manner as previously described [45]. Briefly, 5 × 10^6^ pLB4/11 cells were mixed with 3 μg plasmid DNA (which was first linearized by digestion with NotI) in a total volume of 800 μl PBS at room temperature. The cell/DNA mixture was then electroporated in a 0.4 cm gap electroporation cuvette using a Bio-Rad Gene Pulser (Bio-Rad, Hercules, CA, USA) at 700 V and 25 μF. Following electroporation, cells were plated into medium under no selection for 2 days. Cells were then harvested and plated into 75cm^2^ flasks at a density of 1×10^6^ cells per flask in medium containing 0.5 μg/mL puromycin to select for stable transfectants. After 14 days of selection, colonies that showed nuclear GFP fluorescence were propagated further and GFP-wt lamin A expression was confirmed by Western blot. One clone expressing GFP-wt lamin A was dubbed “pLB4-lamin A” and was used in further experiments.

### Western blotting

Blots were performed using SDS-PAGE with 8% polyacrylamide gels followed by transfer to a nitrocellulose membrane. Each lane contained 20 μg of total cellular protein, and Bio-Rad Precision Plus Kaleidoscope Protein Standard (#161-0375, 5 μL) was used for molecular weight markers. The primary antibody used was GFP (B-2): sc-9996 (mouse monoclonal, from Santa Cruz Biotechnology, Inc.) at a dilution of 1:500. Secondary antibody used was goat anti-mouse IgG-HRP: sc-2005 (Santa Cruz Biotechnology, Inc.) at a dilution of 1:1000. Detection was accomplished using GE Healthcare Amersham ECL Select Western Blotting Detection Reagent and western blot images were analyzed using Typhoon FLA 7000 and ImageQuant LAS 4000 (GE Health).

### Immunofluorescence

Cells (5×10^5^) were seeded onto glass coverslips and grown overnight in six-well plates. The following day, growth medium was removed and cells were washed with 1x PBS. Cells were fixed onto the glass coverslips with 4% formaldehyde in 1x PBS for 10 min at RT. Then, immunofluorescence (IF) was performed as previously described [55]. Briefly, coverslips were blocked for one hour RT in 1-2 mL antibody dilution buffer (2% Bovine Serum Albumin (BSA), 1% Fish Gelatin) in 1x PBS. Then, coverslips were incubated in primary antibody for γH2AX (Bethyl, A300-081A) at a dilution of 1:2000 for one hour at RT. The coverslips were carefully washed three times with PBST for 5 minutes each. Coverslips were incubated with goat anti-rabbit Alexa Fluor 594 secondary antibody (Thermo Fisher Scientific A-11037) at a 1:1000 dilution in antibody dilution buffer for 30 minutes at RT, protected from light. Coverslips were rinsed three times for 3-5 minutes with PBST. All coverslips were dehydrated using an ethanol series (70%, 90%, 100%) for 1-2 minutes, and air-dried protected from light. Coverslips were mounted with Fluoromount-G (Thermo Fisher Scientific 00-4958-02) containing 0.2 μg/mL DAPI (Thermo Fisher Scientific 62248). Images were taken under 40x-60x on an EVOS FL microscope. The number of foci were scored by hand, and at least three coverslips were counted for each cell line. The nuclear signal intensity for γH2AX was also analyzed in CellProfiler and ImageJ. Graphs were plotted using GraphPad Prism software. Nuclear foci signal intensity values were derived from 200-300 nuclei per cell line [55].

In some experiments, after the initial overnight growth of cells on coverslips the growth medium was replaced with medium containing 2 mM HU (MilliporeSigma, 400046). After a two-hour incubation in HU, the media was replaced with fresh growth medium without HU. Cells were either fixed immediately for IF, or were fixed 12hr or 24hr post-HU treatment. IF was performed as described above.

### Cell cycle analysis

Approximately 1 x 10^6^ exponentially growing cells were harvested by trypsinization and pelleted by centrifugation for 5 min. at 200 x g. Cells were then rinsed in one mL ice cold PBS containing 1% BSA and pelleted again. The cell pellet was resuspended in two mL ice cold PBS (without BSA) and fixed by the slow addition of ethanol to a final concentration of 70% at which point the cells were placed at −20°C overnight. The next day, fixative was removed from cells by centrifugation followed by a rinse with 1mL ice-cold PBS with 1% BSA followed by centrifugation. DNA was then labeled by resuspending cells in 1mL of fresh DAPI staining solution (0.1% Triton X-100, 0.1 mg/mL RNase, 1 μg/mL DAPI diluted in 1 x PBS) and run on a BD LSR II Flow Cytometer in the Microscopy and Flow Cytometry Facility at the University of South Carolina, College of Pharmacy. Cell cycle analysis was performed using BD FACS Diva 8.0 software. 30,000 cells were analyzed per experiment.

### Recovery of DSB-induced HR and NHEJ events

In order to induce a DSB at the I-SceI site within the integrated copy of substrate pLB4, cells were electroporated with the I-SceI expression plasmid pCMV-3xnls-I-SceI (“pSce”), generously supplied by Maria Jasin (Sloan Kettering), essentially as previously described [56]. Briefly, 5 × 10^6^ cells were mixed with 20 μg of pSce in a volume of 800 μl of phosphate buffered saline at room temperature and electroporated in a 0.4 cm gap electroporation cuvette using a Bio-Rad Gene Pulser (Bio-Rad, Hercules, CA, USA) set to 700 V and 25 μF. Following electroporation, cells were plated into growth medium under no selection and allowed to recover for 2 days. Cells were then harvested and plated into 75 cm2 flasks at a density of 1 × 10^6^ cells per flask using medium containing 1000 ug/mL of G418 to select for cells that had undergone HR or NHEJ at the DSB site.

### Determination of spontaneous intrachromosomal HR frequencies

Spontaneous recombination frequencies were determined for cell line pLB4-wt lamin A by fluctuation tests that were performed by first generating 10 subclones of the line and then propagating the subclones separately to several million cells each. Cells from each subclone were then plated into several 75 cm^2^ flasks at a density of 3 × 10^6^ cells per flask in order to select for G418^R^ segregants. Colony frequency was then calculated by dividing the number of G418^R^ colonies recovered by the number of cells plated into G418. G418^R^ colonies were harvested for further propagation and analysis.

### PCR amplification and DNA sequence analysis

A segment of the tk-neo fusion gene spanning the I-SceI site was amplified from 500 ng of genomic DNA isolated from G418^R^ clones using primers AW85 (5′- TAATACGACTCACTATAGGGCCAGCGTCTT GTCATTGGCG-3′) and AW91 (5′- GATTTAGGTGACACTATAGCCAAGCGGCCGGAGAACCTG-3′). AW85 is composed of nucleotides 308–327 of the coding sequence of the herpes tk gene, numbering according to [57], with a T7 forward universal primer appended to the 5′ end of the primer. AW91 is composed of 20 nucleotides from the noncoding strand of neomycin gene mapping 25 through 44 bp downstream from the neomycin start codon, with an Sp6 primer appended to the 5′ end of the primer. The positions of the PCR primers are indicated in Fig. 1. PCR was carried out using PuReTaq^TM^ Ready-To-Go^TM^ PCR beads (Cytiva) and a “touchdown” PCR protocol as previously described [58]. PCR products were then sequenced using a T7 primer by Eton Bioscience, Inc. (Research Triangle Park, NC).

**Figure 1.**
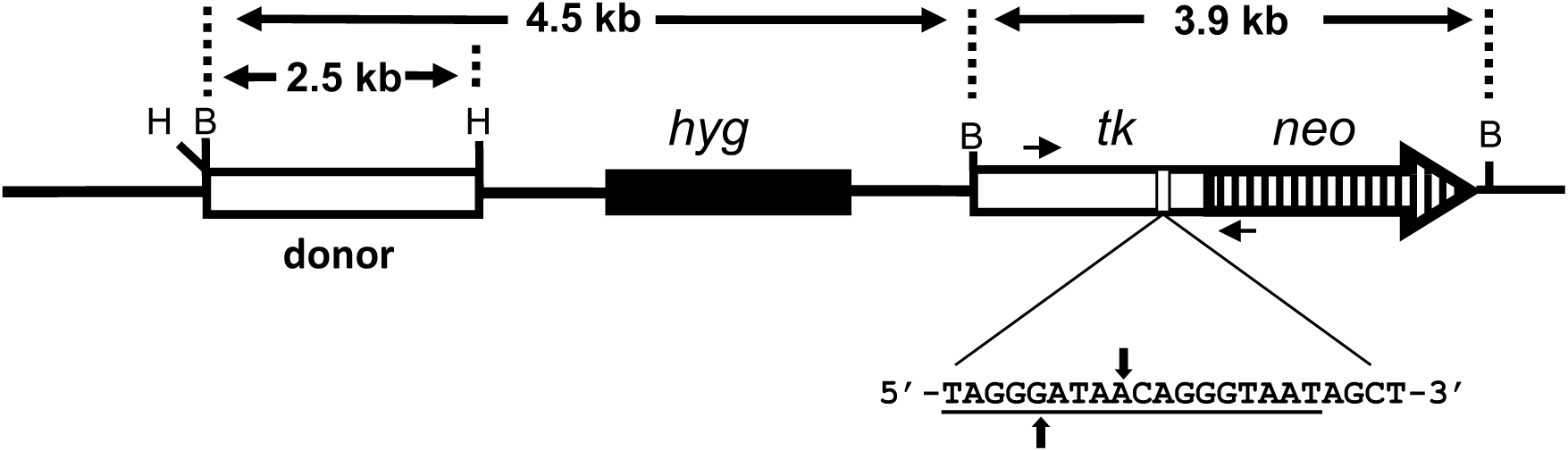
DNA repair substrate pLB4. pLB4 contains a tk-neo fusion gene disrupted by a 22 bp oligonucleotide containing the I-SceI recognition site (underlined). The sites of staggered cleavage are indicated. Short horizontal arrows represent PCR primers used in subsequent analysis of DNA transactions. The donor sequence is a complete functional herpes tk gene. BamHI (B) and HindIII (H) sites are shown.

## Results

### A system to study the impact of progerin on DSB repair in human cells

Our work focuses on gaining a better understanding of the impact that progerin expression has on DSB repair and, more broadly, on genome stability. To do so, we have developed and made use of cell line pLB4/11 which is a derivative of human fibroblast cell line GM637, an SV40-immortalized cell line derived from an apparently healthy individual. Cell line pLB4/11 contains a single stably integrated copy of DNA repair reporter substrate pLB4 (Fig. 1). A DSB can be induced in the I-SceI site within the tk-neo fusion gene on pLB4 by transient expression of I-SceI. Subsequent selection for G418-resistant (G418^R^) clones allows recovery of DSB repair events occurring via NHEJ or HR between the fusion gene and the donor tk sequence.

We previously engineered derivatives of pLB4/11 that constitutively express GFP-progerin and GFP, and we named these cell lines pLB4-progerin and pLB4-GFP, respectively [45]. In the current study we isolated an additional derivative of pLB4/11 that constitutively expresses GFP-wt lamin A and we named this new cell line pLB4-lamin A. The nuclei of pLB4-progerin, but not the other cell lines, are generally misshapen, with some showing blebs (Fig. 2) as seen in HGPS [59].

**Figure 2.**
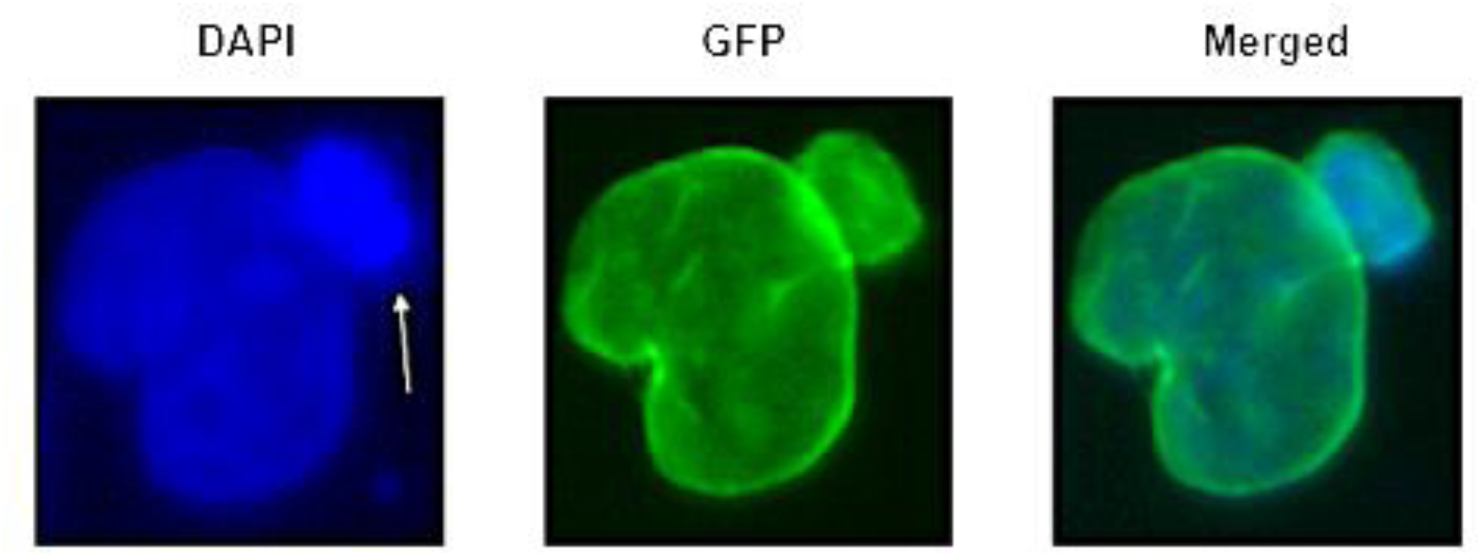
A representative blebbed nucleus from cell line pLB4-progerin. Blebbing is indicated by arrow. DAPI stains DNA and GFP detects GFP-progerin. The bleb appears to contain a high concentration of DNA.

In our previous studies [45] we had shown that the level of expression of GFP-progerin in cell line pLB4-progerin is comparable to the level of progerin expression in a cell line derived from an HGPS patient. As shown by western blot (Fig. 3), expression of GFP-lamin A in cell line pLB4-lamin A is comparable to the expression level of GFP-progerin in line pLB4-progerin.

**Figure 3.**
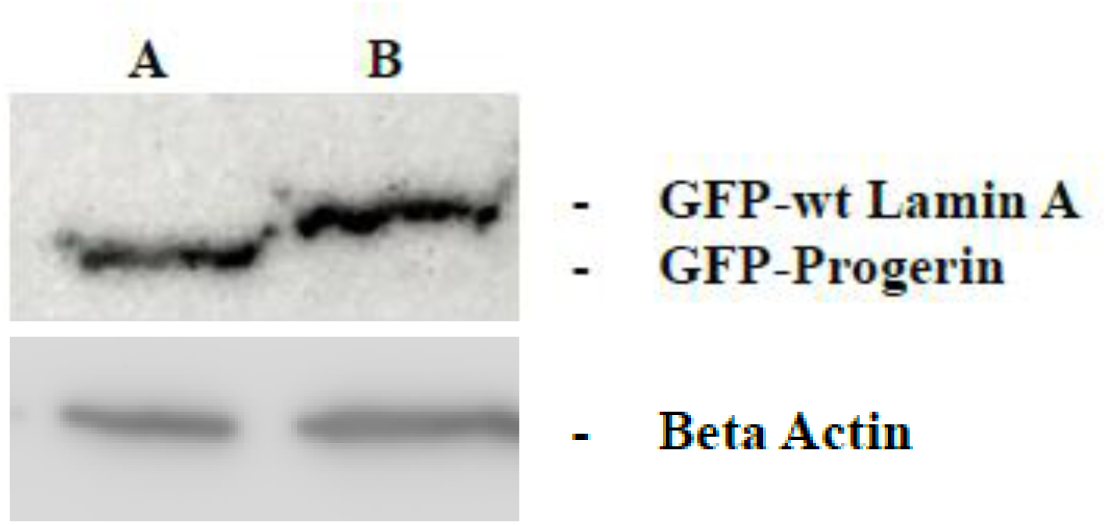
Expression levels of GFP- progerin and GFP- lamin. **A.** Protein from cell line pLB4-progerin is displayed in lane A and protein from cell line pLB4-lamin A is displayed in lane B of a western blot. 30 µg of total cell protein was loaded in each lane. An anti-GFP antibody was used to visualize GFP-progerin and GFP-lamin A. An antibody against beta actin demonstrates equal protein loading in the two lanes.

### Expression of progerin induces endogenous DNA damage in immortalized cells

Comparison of DSB repair in cell line pLB4-progerin with repair in lines pLB4/11 and pLB4-GFP previously revealed that that expression of progerin shifted repair of an I-SceI-induced DSB away from HR and toward NHEJ, and increased the fraction of HR events that had occurred via gene conversion relative to crossing-over [45]. Further, spontaneous HR and gene amplifications were elevated in cells expressing progerin. We were curious to learn if these changes in DNA metabolism were associated with a concomitant increase in the overall level of endogenous DNA damage.

To gain insight into how alterations in nuclear lamina components may impact endogenous genomic damage, we examined the level of spontaneous DNA damage in the genomes of cell lines pLB4/11, pLB4-GFP, pLB4-progerin, and pLB4-lamin A. Our approach was to use a standard immunofluorescence method to score foci of γH2AX, a phosphorylated form of histone H2AX that is produced in the vicinity of DNA damage as an early cellular response to the damage. Damage foci were scored in about 300 nuclei per cell line, and nuclei were binned into categories of nuclei displaying fewer than three damage foci and nuclei with three or more foci. Fig. 4 provides images of γH2AX foci visualized in the cell lines, and quantification of endogenous γH2AX foci in all four cell lines is presented in Table 1A. The data revealed that the portion of nuclei displaying three or more damage foci was significantly greater in cell line pLB4-progerin relative to cell lines pLB4/11, pLB4-GFP, and pLB4-lamin A (p = 2.28 x 10^-8^, p = 5.65 x 10^-4^, and p = 2.69 x 10^-2^, respectively, by chi square). The level of endogenous damage in cell line pLB4-lamin A was elevated relative to damage levels in parent cell line pLB4 (p = 5.61 x 10^-4^) but was not statistically any greater than the damage level seen in pLB4-GFP (p = 0.20). Curiously, pLB4-GFP displayed increased damage relative to pLB4/11 (p = 3 x 10^-2^), suggesting that GFP expression may produce some type of cellular stress. Overall, our results indicated that progerin-expressing cells have elevated levels of endogenous chromosomal damage and that much of this increase is due to expression of progerin rather than simply overexpression of a nuclear lamina component.

**Figure 4:**
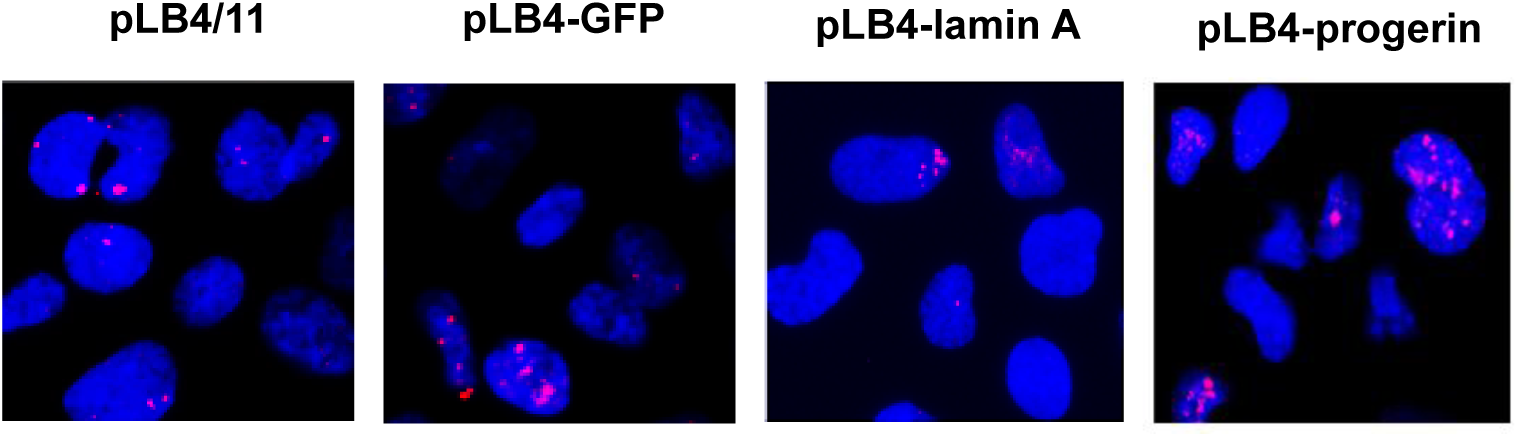
Detection of endogenous DNA damage. Shown are representative images visualizing γH2AX foci in cell lines pLB4/11, pLB4-GFP, pLB4-lamin A, and pLB4-progerin. Cell nuclei were stained blue with DAPI, and immunofluorescence with a Texas-Red conjugated antibody against γH2AX was used to detect damage foci as red dots.

**Table 1.**
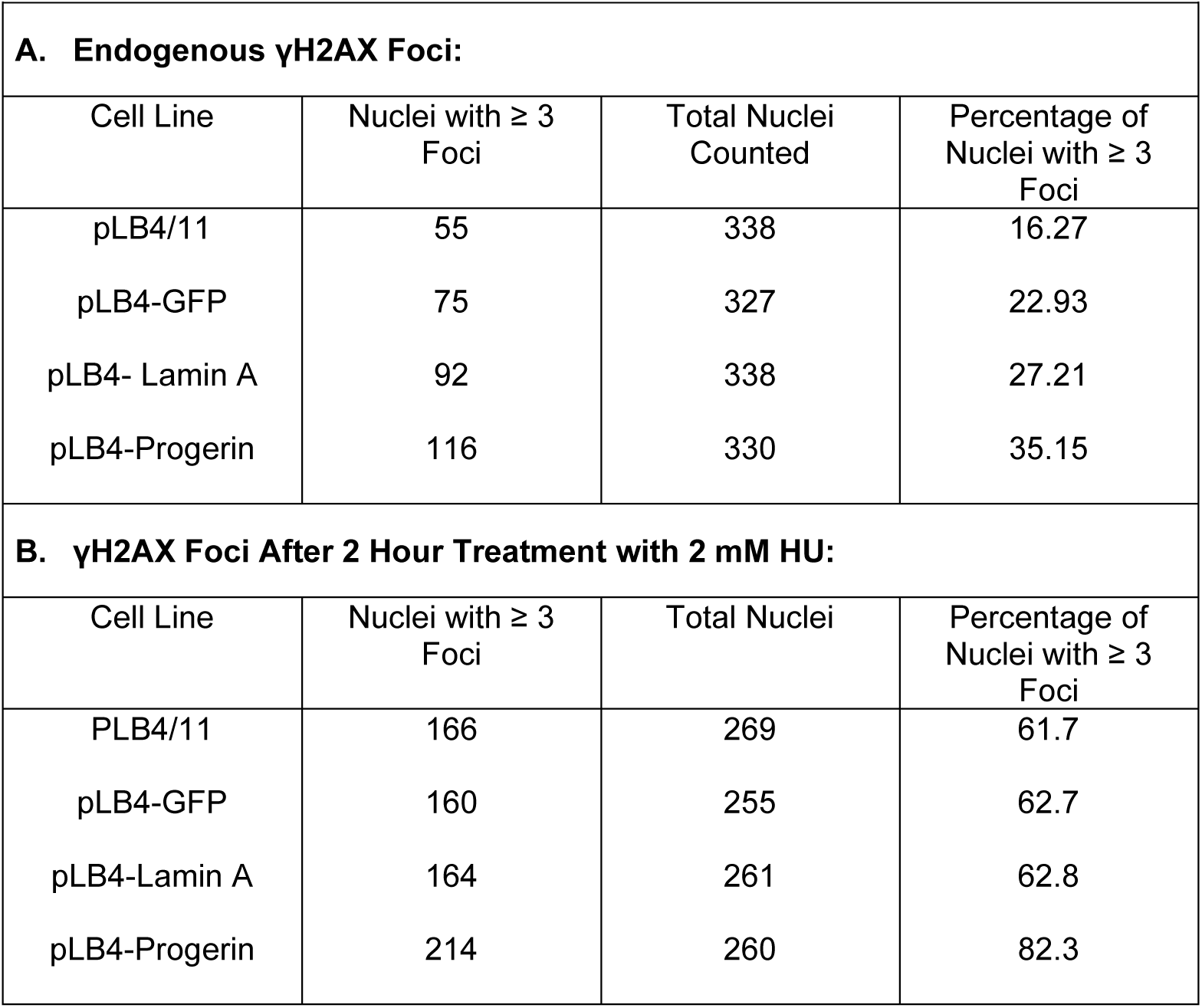
γH2AX Foci Counts.

### Expression of progerin does not alter the cell cycle in immortalized cells

Because progerin-induced DNA damage is often attributed to replication stress in the form of stalled and collapsed replication forks, and because cell line pLB4-progerin accumulated substantially elevated levels of damage relative to it’s parent cell line pLB4/11 (Table 1), we were keen to assess if there was any evidence for replication obstruction in cell line pLB4-progerin. Cell line pLB4-progerin and pLB4/11 are derivatives of normal human fibroblast cell line GM637. We were mindful that GM637 had been immortalized with SV40 and that immortalization disrupts cell cycle checkpoints. Nonetheless, if it were indeed the case that progerin-induced genomic damage is generated primarily as a consequence of a high level of physical disruption of replication fork progression due to impeded loading of replication factors, it stood to reason that we should have observed an alteration in the cell cycle profile in pLB4-progerin cells relative to the parent pLB4/11 line.

As shown in Fig. 5, cell cycle analysis revealed a virtually identical distribution of cells in G1, S, and G2/M for pLB4-progerin and parent cell line pLB4/11. There was no evidence of any cells with sub-G1 DNA content for either cell line, indicating a dearth of apoptotic cells. We concluded that progerin expression in pLB4-progerin did not alter the cell cycle relative to parent line pLB4/11 and did not result in cell death, which indicated that progerin expression did not fundamentally create a barrier to cell proliferation. Further, the doubling time for both pLB4-progerin and parent line pLB4/11 was approximately 24 hours and both cell lines were cultured for several months with no apparent change in doubling time. We therefore concluded that the elevated level of endogenous DNA damage in pLB4-progerin cells was not generated by obstructions to replication fork progression that altered global replication rates.

**Figure 5.**
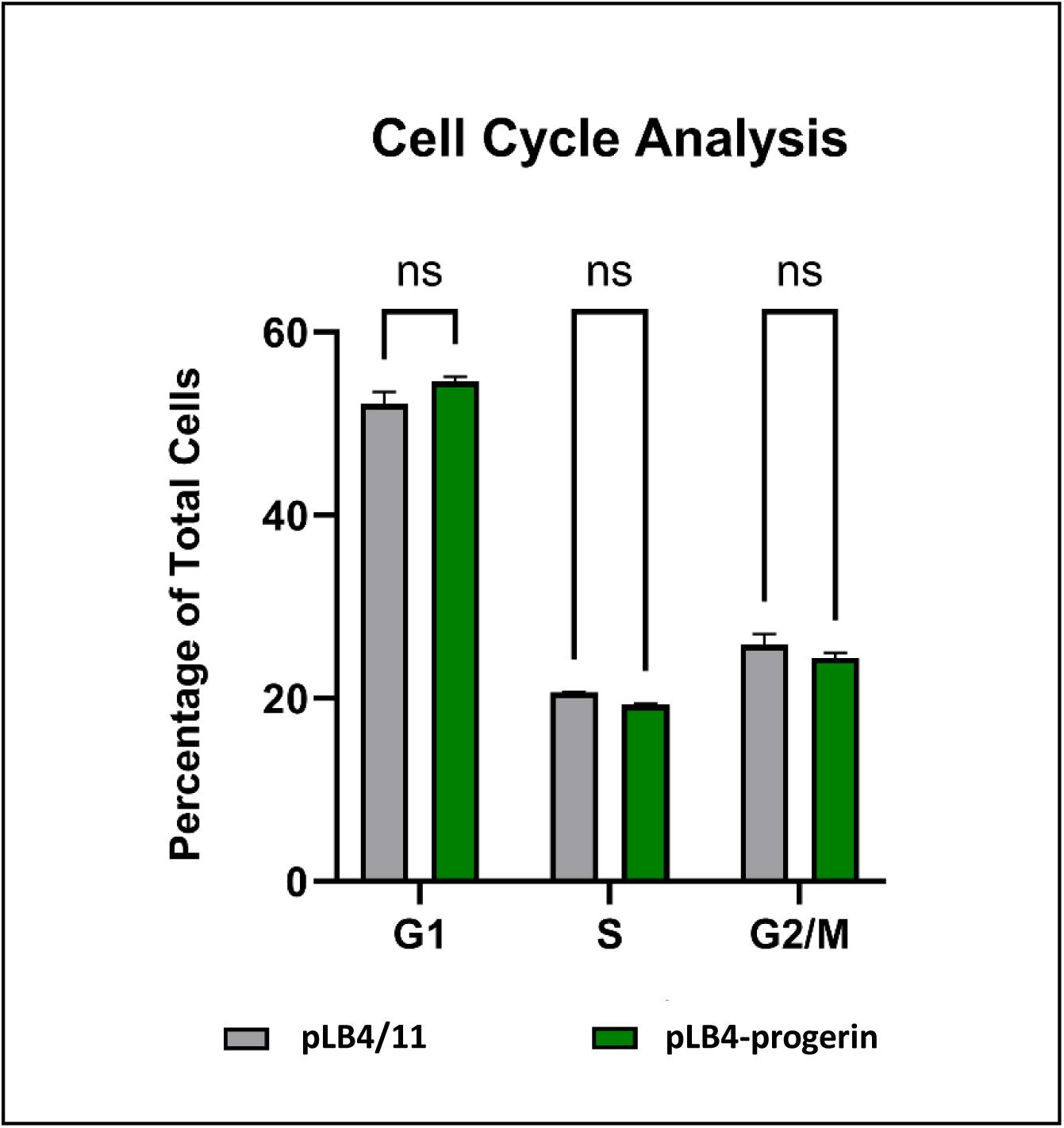
Cell cycle analysis of cell lines pLB4/11 and pLB4-progerin. Shown are the percentages of pLB4/11 and GFP-progerin cells in each cell cycle phase. Two trials were run, with 30,000 cells counted per cell line per trial. Standard deviation is indicated, and there was no significant difference seen between the two cell lines. ns: not significant

### Cells expressing progerin are sensitive to treatment with HU

Our observations indicated that progerin inflicts genomic damage in immortalized cells without causing any obvious impediment to cell proliferation and without imposing any overt obstacle to the progression of replication forks that elicits a cell cycle arrest. Nonetheless, we were curious to learn if exogenous induction of replication fork stalling might induce a particularly high level of damage in cells expressing progerin. To explore this question, we assessed levels of DNA damage following HU treatment of cell lines pLB4/11, pLB4-GFP, pLB4-lamin A, and pLB4-progerin. HU treatment inhibits ribonucleotide reductase and is commonly used to perturb nucleotide pools and bring about subsequent replication fork stalling [60].

After a two-hour treatment with 2 mM HU, all cell lines displayed significantly increased chromosomal damage signaling relative to endogenous damage as measured by the portion of cells displaying three or more γH2AX foci (Table 1B). Cell line pLB4-progerin displayed greater damage after HU treatment than did cell line pLB4/11, pLB4-GFP, or pLB4-lamin A (p = 1.4 x 10^-7^, p = 6.43 x 10^-7^, p = 6.36 x 10^-7^, respectively, by chi square). Strikingly, damage in pLB4-progerin persisted for at least 24 hours post HU-treatment, whereas the level of damage in the three other cell lines had essentially recovered to pre-treatment level by that time point (Fig. 6).

**Fig 6.**
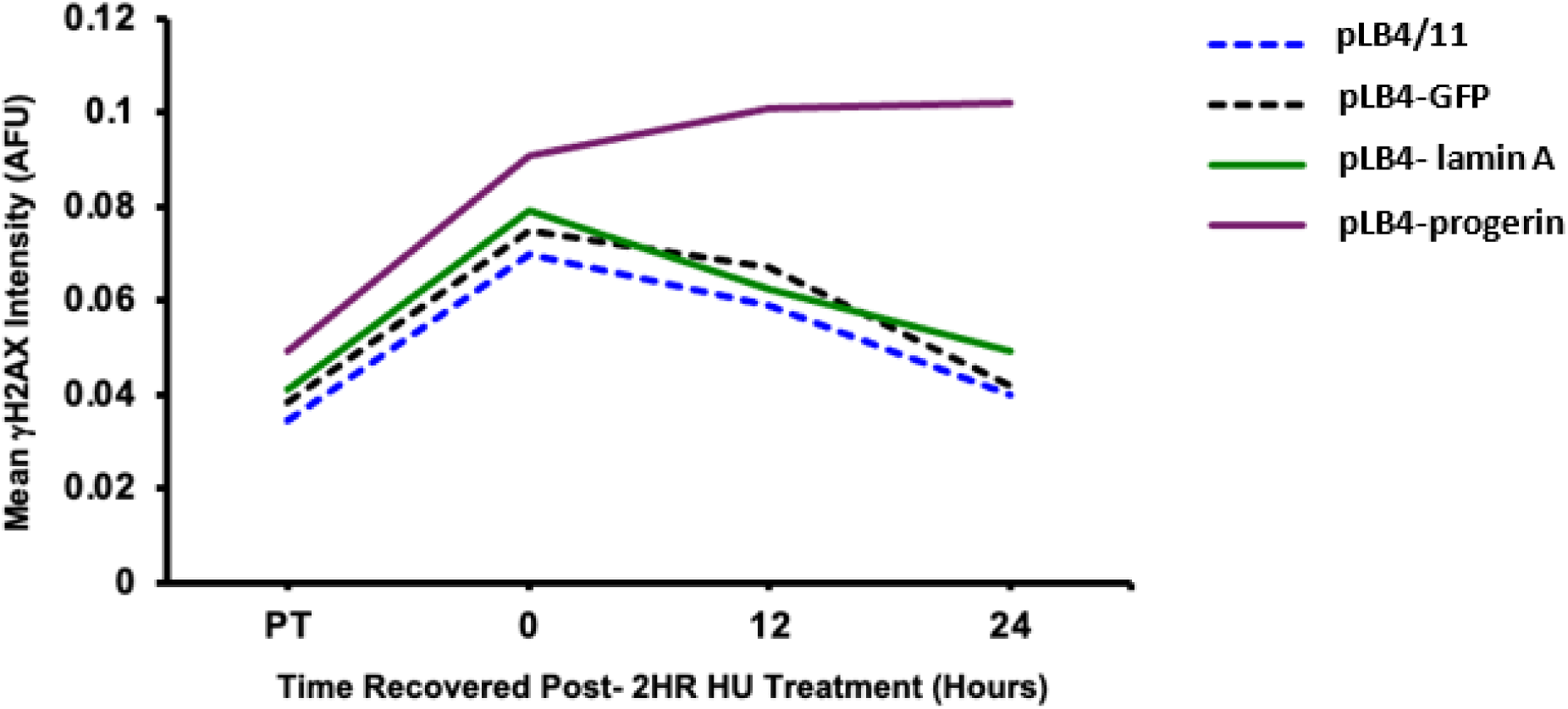
Persistence of DNA damage in cell line GFP-progerin after HU treatment. Shown is a plot of global mean γH2AX immunofluorescence intensity in arbitrary fluorescence units immediately following a 2 hr treatment with 2 mM HU (0 hrs of recovery), and at 12 hrs and 24 hrs of recovery following the HU treatment. PT refers to pre-treatment with γH2AX.

### Overexpression of wt lamin A does not alter genetic recombination and DSB repair

The above experiments demonstrated that cell line pLB4-progerin accumulated significantly more endogenous DNA damage than did pLB4-lamin A, and pLB4-lamin A did not display the sensitivity to HU treatment that pLB4-progerin did. We were curious to further learn whether overexpression of wt lamin A induced any of the alterations to DSB repair and spontaneous recombination that we previously reported for cells expressing progerin.

Cell line pLB4-lamin A was electroporated with plasmid pSce to induce a DSB within the integrated copy of substrate pLB4 and DSB repair events were recovered as G418^R^ colonies. Genomic DNA was isolated from these colonies and DNA sequences surrounding the healed DSB were PCR-amplified. The PCR products were sequenced to determine if each clone arose from HR or NHEJ. HR events are characterized by loss of the 22 bp oligonucleotide that disrupts the fusion gene, and an accompanying transfer of a few scattered heterologous nucleotides from the donor tk gene which serve as markers to identify HR events. Each HR event was further classified as a gene conversion or crossover based on Southern blotting (Fig. 7). Alternatively, DSB repair may occur via NHEJ events which display a deletion, or insertion, at the DSB site that restores the correct reading frame to allow neo expression, with no concomitant transfer of marker nucleotides from the linked functional tk gene. The results of such analyses are summarized in Table 2. The data in table 2 revealed that the shift in DSB repair from HR to NHEJ observed for cell line pLB4-progerin (relative to DSB repair events from pLB4/11 and pLB4-GFP) was not observed for cell line pLB4-lamin A.

**Figure 7.**
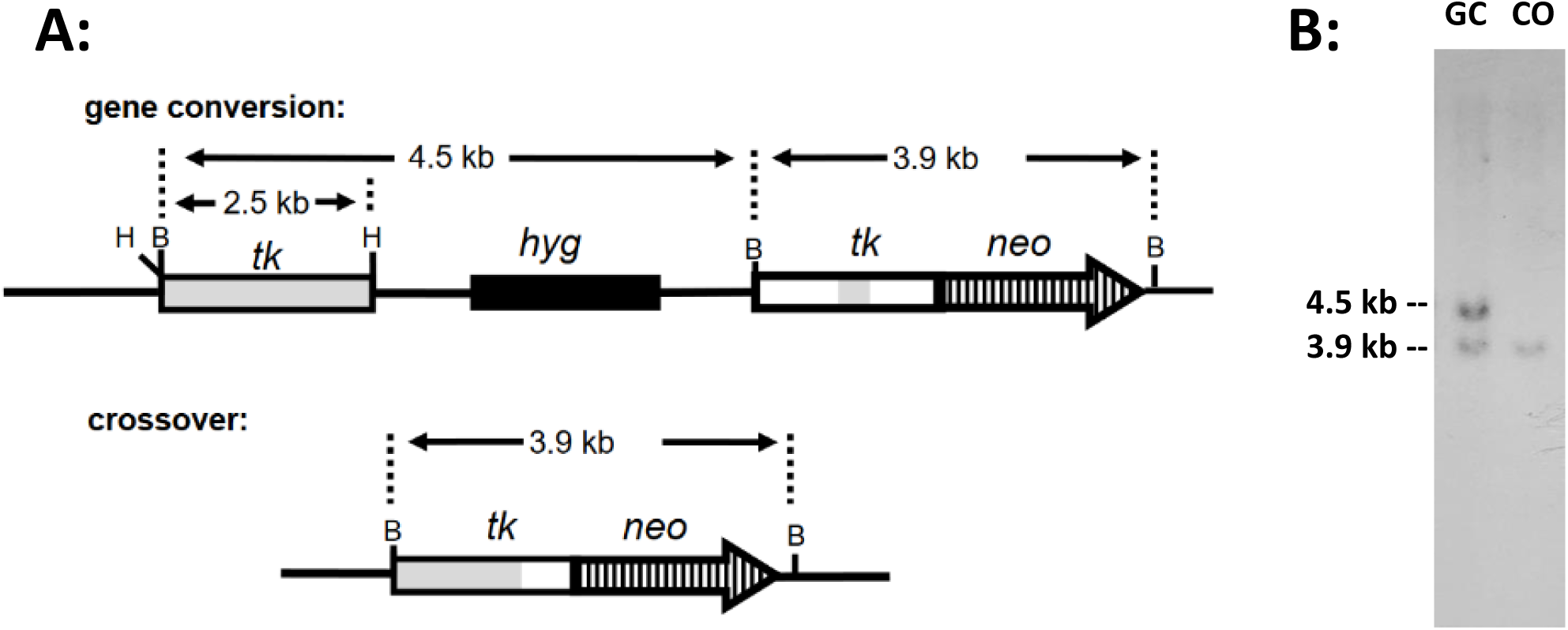
Distinguishing gene conversions from crossovers. Panel A-- Schematic showing that a BamHI digestion of genomic DNA isolated from a gene conversion clone produces a 4.5 kb and a 3.9 kb fragment predicted to be detectable on a Southern blot using a tk probe. In contrast, a crossover clone produces only a 3.9 kb fragment. Panel B-- Representative Southern blot using a tk probe to display BamHI digestions of DNA isolated from a clone that arose from a gene conversion (GC) and a clone that arose from a crossover (CO).

**Table 2.**
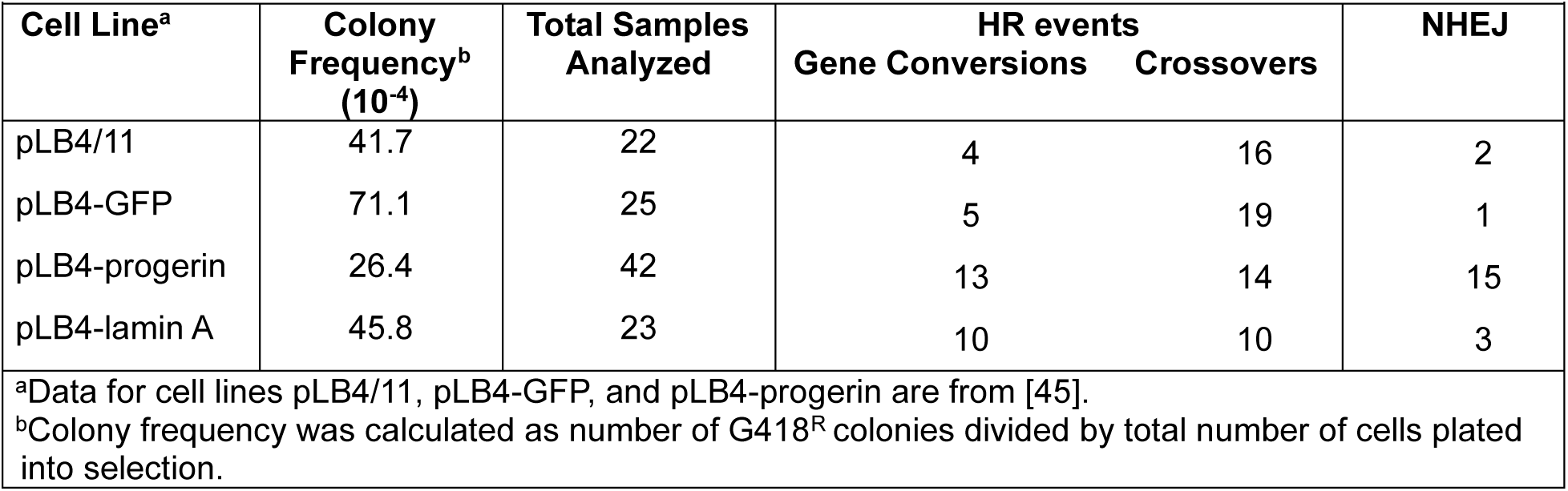
Analysis of recovered DSB repair events.

| Table 2. Analysis of recovered DSB repair events |  |  |  |  |  |
| --- | --- | --- | --- | --- | --- |
| Cell Line<sup>a</sup> | Colony Frequency<sup>b</sup><br>(10<sup>-4</sup>) | Total Samples Analyzed | HR events |  | NHEJ |
|  |  |  | Gene Conversions | Crossovers |  |
| pLB4/11 | 41.7 | 22 | 4 | 16 | 2 |
| pLB4-GFP | 71.1 | 25 | 5 | 19 | 1 |
| pLB4-progerin | 26.4 | 42 | 13 | 14 | 15 |
| pLB4-lamin A | 45.8 | 23 | 10 | 10 | 3 |

<sup>a</sup>Data for cell lines pLB4/11, pLB4-GFP, and pLB4-progerin are from [45].
<sup>b</sup>Colony frequency was calculated as number of G418<sup>R</sup> colonies divided by total number of cells plated into selection.

We also carried out two fluctuation tests on pLB4-lamin A to measure the frequency of formation of spontaneous G418^R^ segregants (Table 3). As presented in Table 3, the frequency with which G418^R^ segregants arose in cell line pLB4-lamin A was no greater than the frequency of G418^R^ segregants in cell lines pLB4/11 and pLB4-GFP. Thus, pLB4-lamin A differed from cell line pLB4-progerin which displayed a substantially elevated frequency of G418 segregants as we previously reported [45]. Further, every analyzed G418^R^ clone that arose from pLB4-lamin A represented a bona-fide HR event (Table 4). This is significantly different (p = 3.6 x 10-3 by a two-sided Fisher exact test) from the 19 out of 35 G418^R^ clones from pLB4-progerin which were previously found to have arisen through a mechanism other than bona-fide HR and appeared to involve a gene amplification event [45]. Additionally, the exclusive recovery of gene conversions among the HR events occurring in pLB4-progerin was not repeated for pLB4-lamin A. In summary, overexpression of wt lamin A did not elicit the alterations in DSB repair and spontaneous genetic rearrangements displayed by cells expressing progerin.

**Table 3.** Recovery of spontaneous G418 resistant segregants.

| **Table 3. Recovery of spontaneous G418 resistant segregants** |  |  |  |  |  |
| --- | --- | --- | --- | --- | --- |
| **Cell Line<sup>a</sup>** | **Cells Plated (10<sup>6</sup>)<sup>b</sup>** | **G418<sup>R</sup> Colonies<sup>c</sup>** | **Median No. Colonies<sup>d</sup>** | **Colony Frequency (10<sup>7</sup>)<sup>e</sup>** | **HR Frequency (10<sup>-7</sup>)<sup>f</sup>** |
| pLB4/11 | 145 (30) | 29 | 0 | 2.0 | 1.8 |
| pLB4-GFP | 30 (10) | 20 | 2 | 6.8 | 6.8 |
| pLB4-progerin | 30 (10) | 228 | 17.5 | 76 | 34.7 |
| pLB4-lamin A | 65 (20) | 64 | 1 | 9.8 (3.9)<sup>g</sup> | 9.8 (3.9)<sup>g</sup> |

<sup>a</sup>Data for cell lines pLB4/11, pLB4-GFP, and pLB4-progerin are from our previous study [45]. <sup>b</sup>For each cell line, 10 or more subclones were each propagated separately to a total of 3–5 million cells before being plated into medium containing G418. Total number of subclones plated is presented in parentheses. <sup>c</sup>Total number of G418<sup>R</sup> colonies recovered from all of the subclones for each cell line. <sup>d</sup>Median number of colonies recovered from all of the subclones plated. <sup>e</sup>Total number of colonies recovered divided by the total number of cells plated. <sup>f</sup>Calculated as colony frequency multiplied by percentage of colonies characterized as bona fide HR. <sup>g</sup>pLB4-lamin A included a single “jackpot” subclone that produced 40 out of the total of 64 colonies recovered. The number in parentheses is the colony frequency and HR frequency for pLB4-lamin A excluding the jackpot subclone.

**Table 4.** Analysis of spontaneous G418^R^ segregants.

| **Table 4. Analysis of spontaneous G418<sup>R</sup> segregants** |  |  |  |  |
| --- | --- | --- | --- | --- |
| **Cell Line<sup>a</sup>** | **No. Clones Analyzed** | **Gene Conversions** | **Crossovers** | **Gene Amplification<sup>b</sup>** |
| pLB4/11 | 24 | 5 | 16 | 3 |
| pLB4-GFP | 17 | 8 | 9 | 0 |
| pLB4-progerin | 35 | 16 | 0 | 19 |
| pLB4-lamin A | 16 | 12 | 5 | 0 |

<sup>a</sup>Analyses of clones from cell lines pLB4/11, pLB4-GFP, and pLB4-progerin are from our previous study [45]. <sup>b</sup>Clones displaying amplification of the number copies of integrated pLB4 substrate with no other sequence change. These clones were recovered due to apparent leaky expression of the parent tk-neo fusion gene which provided G418 resistance when present in multiple copies.

## Discussion

We report here that expression of progerin is associated with a significant increase in the accumulation of DNA damage in an SV40-immortalized human cell line. We recorded elevated levels of endogenous DNA damage in cell line pLB4-progerin which expresses progerin but shows no alteration in cell cycle profile or doubling time relative to its parent cell line pLB4/11 which does not express detectable levels of progerin [45]. Our work demonstrates that progerin expression can inflict damage on the human genome in a manner independent from global replication fork stalling and collapse.

Progerin-induced DNA damage was previously reported for a human fibroblast cell line that had been immortalized by human telomerase reverse transcriptase (hTERT) [61], and our current study demonstrates that progerin’s ability to trigger damage in immortalized cells is not restricted to cells immortalized by hTERT. Moreover, the relationship between hTERT expression and progerin is complicated with regards to both progerin’s effect on DNA damage as well as hTERT’s ability to immortalize cells. Several reports found that hTERT expression protects against progerin-induced DNA damage [62,63], while at least one report found that cells from HGPS patients frequently fail to immortalize with hTERT expression [64].

Our basic observation that progerin expression caused DNA damage in a cell line despite immortalization is relevant to understanding the temporal sequence of events that produces accelerated or normal aging. A commonly accepted paradigm to explain progerin-induced damage posits that progerin reduces replicative capacity by interfering with the recruitment of replication proteins, including PCNA, that are required for replication fork progression. In this scenario, replication forks in progerin-expressing cells are subject to stalling and collapse, leading to strand breakage [33–36, 47–51]. At the same time, nucleotide excision protein XPA is inappropriately recruited to the stalled or collapsed replication forks due to an affinity of XPA for DNA structures that contain a junction between single and double-stranded DNA. Binding of XPA at the replication forks is presumed to be facilitated by exposed DNA which would normally be protected by bound PCNA. The presence of XPA results in the stearic blockage of repair proteins and so DSBs accumulate at the stalled and collapsed forks [34,35,48]. However, since there is no evidence that the progression of replication forks is impeded at a global level in immortalized cell line pLB4-progerin in our study, the stalled replication fork paradigm for the genesis of damage does not provide a satisfactory explanation for the endogenous damage seen in pLB4-progerin.

If replication fork stalling is not a cause of the elevated levels of endogenous damage seen in pLB4-progerin, then what is? The genesis of the accumulated damage remains unknown at this point, but we recognize that in cell line pLB4-progerin there may be either an elevated rate of damage production, a reduced rate of repair, or both. In any case, we infer that essentially all DNA damage in most pLB4-progerin cells is repaired within the course of a cell cycle, since if this were not the case we would expect that the pLB4-progerin cell line would display a notable defect in propagation. Said differently, it is difficult to imagine a cell line successfully proliferating for months with no appreciable cell death if its genome is continually accruing unrepaired damage. We also note that mitotic chromosome spreads of pLB4-progerin did not reveal any abnormal chromosomal structures (not shown).

Although our data points to progerin-inflicted DNA damage occurring independently from impediment of replication fork progression, our experiments did reveal a sensitivity of pLB4-progerin cells to treatment with HU. HU blocks ribonucleotide reductase and is known to induce replication fork stalling and nucleotide pool disruption [60]. The substantially elevated level of damage seen in cell line pLB4-progerin upon HU treatment, along with the persistence of the damage specifically in pLB4-progerin (Fig. 6), implies that the mechanisms for rescuing stalled or collapsed replication forks are impaired by progerin. Pathways that underpin replication as means for rescuing stalled or collapsed forks are believed to involve processes such as DNA strand invasion and annealing of homologous DNA sequences, branch migration, and formation and resolution of Holliday junctions [60]. These processes are in essence equivalent to HR, and replication fork restart indeed often requires the involvement of HR proteins including Rad51 [60]. HR processes are precisely the types of repair processes that we and others have found to be attenuated in cells expressing progerin [41–46], providing an explanation for why stalled replication forks, when they occur, present a particular challenge for progerin-expressing cells to overcome.

It is also a distinct possibility that pLB4-progerin cells are especially sensitive to the diminution of nucleotide pools which is a consequence of HU treatment. Consistent with the notion of progerin-induced sensitivity to nucleotide pool levels, it has been reported that the replication timing signature is perturbed in cells expressing progerin, creating a situation where too many replication origins are active at once [63]. Simultaneous replication from many origins can render cells prone to nucleotide pool exhaustion and can limit the replicative capacity of cells expressing progerin [63]. Significantly, replicative capacity was shown to be restored upon nucleotide supplementation of growth medium [63]. In our current work, there is no evidence that pLB4-progerin displays reduced replicative capacity. Instead, we conjecture that the increased load of endogenous DNA damage observed in cell line pLB4-progerin may place a strain on nucleotide pools since DNA repair either needs to keep up with an elevated rate of production of DNA lesions or needs to play “catch-up” to eliminate damage that, due to possibly sluggish repair, accumulates prior to replication. Progerin-expressing cells are thus effectively faced with repairing many lesions simultaneously in order to complete a cell cycle. The picture painted is that cells expressing progerin may be teetering on the edge of sufficient levels of nucleotides needed to both complete repair and replicate the genome. Interestingly, insufficient levels of purine nucleotides due to reduced levels of ribose-phosphate pyrophosphokinase has been reported as a potentially important contributing factor to premature aging in HGPS [65]. Furthermore, increased ribonucleotide reductase activity has been shown to alleviate the severity of a progeroid disease in mice [66].

In our previous work [45], we reported that the spontaneous rate of HR in pLB4-progerin was elevated relative to cell lines pLB4/11 and pLB4-GFP. We speculated that the elevated frequency of HR in pLB4-progerin may be provoked by high levels of genomic DSBs, despite repair of any given DSB being shifted toward NHEJ. We did not measure genomic damage in our earlier work, but our current work corroborates that pLB4-progerin does display elevated endogenous damage, lending credence to our prior speculation. Our current work also shows that overexpression of wt lamin A in pLB4-lamin A does not lead to the level of endogenous damage seen in pLB4-progerin (Table 1), and the frequency of spontaneous HR in pLB4-lamin A is correspondingly not elevated relative to pLB4/11 or pLB4-GFP (Table 3). Further, we do not see evidence of elevated levels of gene amplification in pLB4-lamin A (Table 4) nor do we see a shift in DSB repair pathways from HR to NHEJ in pLB4-lamin A. In short, our results indicate the changes we see in pLB4-progerin are progerin-specific and are not duplicated by disrupting the stoichiometry of nuclear lamina components by overexpressing lamin A.

In summary, our work illustrates that progerin expression can produce elevated levels of endogenous DNA damage independent from an impact on replication progress and cell proliferation. Although replication fork stalling is not required in order for progerin to induce DNA damage, cells expressing progerin appear to have difficulty recovering from damage inflicted by replication fork stalling should it occur. Our work, in conjunction with studies by others, raises the possibility that progerin may force cells to live on the edge when it comes to nucleotide pools in the sense that nucleotides may frequently become a critically limiting commodity. Limiting nucleotides may contribute to a cascade of events producing alterations in DNA repair pathways, accumulation of DNA damage, and obstruction of replication. The interrelatedness between these various aspects of nucleic acid metabolism may set up a vicious cycle rather than a linear temporal sequence of events that leads to cellular demise that contributes to both accelerated and normal aging. By using pLB4-progerin which is an immortalized cell line, obstructed replication was not a contributor to the impact of progerin observed in our studies. It will be instructive to learn if nucleotide supplementation or other treatments of cell line pLB4-progerin can reverse the DNA repair and/or DNA damage phenotypes recorded for this progerin-expressing cell line. Such investigations using our model system can potentially allow further dissection of the individual elements of the genome instability cycle set in motion by progerin, with the goal of someday learning how to break that cycle.

## Acknowledgements

This work was supported by the National Institute on Aging grant R03AG064525 and the University of South Carolina ASPIRE Proposal 130100-22-60150 to A.S.W., and a University of South Carolina Magellan Scholar award to T.M.V.

## Declaration of Competing Interest

The authors report there are no competing interests to declare.

## Data availability statement

Cell lines and plasmids are available upon request. The authors affirm that all data necessary for confirming the conclusions of the article are present within the article, figures, and tables.

## Notes

### Competing Interest Statement

The authors have declared no competing interest.

## References

[1] E. Sonoda, H. Hochegger, A. Saberi, Y. Taniguchi, S. Takeda, Differential usage of non-homologous end-joining and homologous recombination in double-strand break repair, DNA Repair 5 (2006) 1021–1029.

[2] J. San Filippo, P. Sung, H. Klein, Mechanism of eukaryotic homologous recombination, Annu. Rev. Biochem. 77 (2008) 229–257.

[3] B. Pardo, B. G‘omez-Gonzalez, A. Aguilera, DNA double strand break repair: how to fix a broken relationship, Cell. Mol. Life Sci. 66 (2009) 1039–1056.

[4] M. Shrivastav, L.P. De Haro, J.A. Nickoloff, Regulation of DNA double-strand break repair pathway choice, Cell Res. 18 (2009) 134–147.

[5] W.D. Heyer, K.T. Ehmsen, J. Liu, Regulation of homologous recombination in eukaryotes, Annu. Rev. Genet. 44 (2010) 113–139.

[6] E.M. Kass, M. Jasin, Collaboration and competition between DNA double-strand break repair pathways, FEBS Lett. 584 (2010) 3703–3708.

[7] J.R. Chapman, M.R. Taylor, S.J. Boulton, Playing the end game: DNA double-strand break repair pathway choice, Mol. Cell 47 (2012) 497–510, 10.1016/j.molcel.2012.07.029.

[8] A.A. Goodarzi, P.A. Jeggo, The repair and signaling responses to DNA double-strand breaks, Adv. Genet. 82 (2013) 1–45, 10.1016/B978-0-12-407676-1.00001-9.

[9] T. Iyama, D.M. Wilson III, DNA repair mechanisms in dividing and non-dividing cells, DNA Repair 12 (2013) 620–636, 10.1016/j.dnarep.2013.04.015.

[10] R. Scully, A. Panday, R. Elango, N.A. Willis, DNA double-strand break repair pathway choice in somatic mammalian cells, Nat. Rev. Mol. Cell Biol. 20 (2019) 698–714, 10.1038/s41580-019-0152-0.

[11] A.J. Bishop, R.H. Schiestl, Role of homologous recombination in carcinogenesis, Exp. Mol. Pathol. 74 (2003) 94–105.

[12] J.E. Eyfjord, S.K. Bodvarsdottir, Genomic instability and cancer: networks involved in response to DNA damage, Mutat. Res. 592 (2005) 18–28.

[13] R.D. Kennedy, A.D. D’Andrea, DNA repair pathways in clinical practice: lessons from pediatric cancer susceptibility syndromes, J. Clin. Oncol. 24 (2006) 3799–3808.

[14] V. Gorbunova, A. Seluanov, A, Making ends meet in old age: DSB repair and aging, Mech. Ageing Dev. 126 (2005) 621–628.

[15] K.J. Kyng, V.A. Bohr, Gene expression and DNA repair in progeroid syndromes and human aging, Ageing Res. Rev. 4 (2005) 570–602.

[16] D.B. Lombard, K.F. Chua, R. Mostoslavsky, S. Franco, M. Gostissa, F.W. Alt, DNA repair, genome stability, and aging, Cell 120 (2005) 497–512.

[17] V. Gorbunova, A. Seluanov, Z. Mao, C. Hine, Changes in DNA repair during aging, Nucleic Acids Res. 35 (2007) 7466–7474.

[18] A.A. Freitas, J.P. de Magalhaes, A review and appraisal of the DNA damage theory of ageing, Mutat. Res. 728 (2011) 12–22.

[19] V. Tiwari, D.M. Wilson 3rd, DNA damage and associated DNA repair defects in disease and premature aging, Am. J. Hum. Genet. 105 (2019) 237–257, 10.1016/j.ajhg.2019.06.005.

[20] K. Ren, S.P. de Ortiz, Non-homologous DNA end joining in the mature rat brain, J. Neurochem. 80 (2002) 949–959.

[21] J. Vijg, M.E.T. Dolle, Large genome rearrangements as a primary cause of aging, Mech. Ageing Dev. 123 (2002) 907–915.

[22] A. Seluanov, D. Mittelman, O.M. Pereira-Smith, J.H. Wilson, V. Gorbunova, DNA end joining becomes less efficient and more error-prone during cellular senescence, Proc. Natl. Acad. Sci. USA 101 (2004) 7624–7629.

[23] C.Z. Bachrati, I.D. Hickson, RecQ helicases: suppressors of tumorigenesis and premature aging, Biochem. J. 374 (2003) 577–606.

[24] M. O’Driscoll, P.A. Jeggo, The role of double-strand break repair - insights from human genetics, Nat. Rev. Genet. 7 (2006) 45–54.

[25] R.M. Brosh Jr., V.A. Bohr, Human premature aging, DNA repair and RecQ Helicases, Nucleic Acids Res. 35 (2007) 7527–7544.

[26] C. Ĺopez-Otín, M.A. Blasco, L. Partridge, M. Serrano, G. Kroemer, The hallmarks of aging, Cell 153 (2013) 1194–1217, 10.1016/j.cell.2013.05.039.

[27] R. Burla, M. La Torre, C. Merigliano, F. Vernì, I. Saggio, Genomic instability and DNA replication defects in progeroid syndromes, Nucleus 9 (2018) 368–379, 10.1080/19491034.2018.1476793.

[28] M.S. Ahmed, S. Ikram, N. Bibi, A. Mir, Hutchinson-Gilford progeria syndrome: a premature aging disease, Mol. Neurobiol. 55 (2018) 4417–4427, 10.1007/s12035-017-0610-7.

[29] K. Cao, C.D. Blair, D.A. Faddah, J.E. Kieckhaefer, M. Olive, M.R. Erdos, E.G. Nabel, F.S. Collins, Progerin and telomere dysfunction collaborate to trigger cellular senescence in normal human fibroblasts, J. Clin. Invest. 121 (2011) 2833–2844.

[30] D. McClintock, D. Ratner, M. Lokuge, D.M. Owens, L.B. Gordon, F.S. Collins, K. Djabali, The mutant form of lamin A that causes Hutchinson-Gilford Progeria is a biomarker of cellular aging in human skin, PLoS One 2 (2007), e1269, 10.1371/journal.pone.0001269.

[31] P. Scaffidi, T. Misteli, Lamin A-dependent nuclear defects in human aging, Science 312 (2006) 1059–1063.

[32] V.V. Ashapkin, L.I. Kutueva, S.Y. Kurchashova, I.I. Kireev, Are there common mechanisms between the Hutchinson-Gilford Progeria Syndrome and Natural Aging? Front. Genet. 10 (2019) 455, 10.3389/fgene.2019.00455.

[33] R.N. Serio, Unraveling the mysteries of aging through a Hutchinson–Gilford progeria syndrome model, Rejuvenation Res. 14 (2011)133–141.

[34] P.R. Musich and Y. Zou, Genomic instability and DNA damage responses in progeria arising from defective maturation of prelamin A, Aging 1 (2009) 28–37.

[35] P.R. Musich and Y. Zou, DNA-damage accumulation and replicative arrest in Hutchinson–Gilford progeria syndrome, Biochemical Soc. Trans. 39 (2011)1764–1769.

[36] Y. Liu, Y. Wang, A.E. Rusinol, M.S. Sinensky, J. Liu, J, S.M. Shell, and Y. Zou, Involvement of xeroderma pigmentosum group A (XPA) in progeria arising from defective maturation of prelamin A, FASEB J. 22 (2008) 603–611.

[37] D. Constantinescu, A.B. Csoka, C.S. Navara, and G.P. Schatten, Defective DSB repair correlates with abnormal nuclear morphology and is improved with FTI treatment in Hutchinson-Gilford progeria syndrome fibroblasts, Exp. Cell Res. 316 (2010) 2747–2759.

[38] S.A. Richards, J. Muter, P. Ritchie, G. Lattanzi, and C.J. Hutchison, The accumulation of un-repairable DNA damage in laminopathy progeria fibroblasts is caused by ROS generation and is prevented by treatment with N-acetyl cysteine, Hum. Mol. Genet. 20 (2011)3997–4004.

[39] C.J. Hutchison, The role of DNA damage in laminopathy progeroid syndromes, Biochem. Soc. Trans. 39 (2011) 1715–1718.

[40] I. Gonzalez-Suarez, and S. Gonzalo, Nurturing the genome: A-type lamins preserve genomic stability, Nucleus 1 (2010) 129–135

[41] S. Gonzalo, DNA damage and lamins, Adv. Exp. Med. Biol. 773 (2014)377–399.

[42] H. Zhang, Z.M. Xiong, and K. Cao, Mechanisms controlling the smooth muscle cell death in progeria via down-regulation of poly(ADP-ribose) polymerase 1, Proc. Natl. Acad. Sci. USA 111 (2014) E2261–E2270.

[43] B. Liu, J. Wang, K.M. Chan, W.M. Tjia, W Deng, X. Guan, J.D. Huang, K.M., P.Y. Chau, D.J. Chen, D. Pei, A.M. Pendas, J. Cadinanos, C. Lopez-Otin, H.F. Tse, C. Hutchison, J. Chen, Y. Cao, K.S. Cheah, K. Tryggvason, and Z. Zhou, Genomic instability in laminopathy-based premature aging, Nat. Med. 11 (2005) 780–785.

[44] S. Gonzalo, and R. Kreienkamp, (2015) DNA repair defects and genome instability in Hutchinson-Gilford Progeria Syndrome, Curr. Opin. Cell. Biol. 34 (2015) 75–83, 10.1016/j.ceb.2015.05.007.

[45] C.J. Komari, A.O. Guttman, S.R. Carr, T. Trachtenberg, E.A. Orloff, A.V. Haas, A.R. Patrick, S. Chowdhary, B.C. Waldman, and A.S. Waldman, Alteration of genetic recombination and double-strand break repair in human cells by progerin expression, DNA Repair 96 (2020) 102975, 10.1016/j.dnarep.2020.102975

[46] L.A. Joudeh, A.J. DiCintio, M.R. Ries, A.S. Gasperson, K.E. Griffin, V.P. Robbins, M. Bonner, S. Nolan, E. Black, and A.S. Waldman, Corruption of DNA end-joining in mammalian chromosomes by progerin expression, DNA Repair 126 (2023) 103491, 10.1016/j.dnarep.2023.103491.

[47] O. Dreesen, Towards delineating the chain of events that cause premature senescence in the accelerated aging syndrome Hutchinson–Gilford progeria (HGPS), Biochemical Society Transactions 48(2020) 981–991, 10.1042/BST20190882

[48] B.A. Hilton, J. Liu, B.M Cartwright, Y. Liu, M. Breitman, Y. Wang, et al., Progerin sequestration of PCNA promotes replication fork collapse and mislocalization of XPA in laminopathy-related progeroid syndromes, FASEB J. 31 (2017) 3882–3893, 10.1096/fj.201700014R

[49] R. Burla, M. La Torre, C. Meriglianoa, F. Vernì, and I.abella Saggio, Genomic instability and DNA replication defects in progeroid syndromes, Nucleus 9 (2018) 368–379, 10.1080/19491034.2018.1476793

[50] R. Kreienkamp, S. Graziano, N. Coll-Bonfill, et al., A cell-intrinsic interferon-like response links replication stress to cellular aging caused by progerin, Cell Reports 22 (2018) 2006–2015, 10.1016/j.celrep.2018.01.090

[51] K. Wheaton, D. Campuzano, M. Weili, M. Sheinis, B. Ho, G.W. Brown, et al., Progerin-induced replication stress facilitates premature senescence in Hutchinson– Gilford progeria syndrome. Mol. Cell. Biol. 37 (2017) 1–18 10.1128/MCB.00659-16

[52] J.A. Smith, L.A. Bannister, V. Bhattacharjee, Y. Wang, B.C. Waldman, A.S. Waldman, Accurate homologous recombination is a prominent double-strand break repair pathway in mammalian chromosomes and is modulated by mismatch repair protein Msh2, Mol. Cell. Biol. 27 (2007) 7816–7827.

[53] L.A. Bannister, B.C. Waldman, A.S. Waldman, Modulation of error-prone double-strand break repair in mammalian chromosomes by DNA mismatch repair protein Mlh1, DNA Repair 3 (2004) 465–474.

[54] B.C. Waldman, Y. Wang, K. Kilaru, Z. Yang, A. Bhasin, M.D. Wyatt, A.S. Waldman, Induction of intrachromosomal homologous recombination in human cells by raltitrexed, an inhibitor of thymidylate synthase, DNA Repair 7 (2008) 1624–1635

[55] Y. Wang, K.S. Brady, B.P. Caiello, S.M. Ackerson, and J.A. Stewart, Human CST suppresses origin licensing and promotes AND-1/Ctf4 chromatin association, Life Sci Alliance 2 (2019) e201800270, 10.26508/lsa.201800270.

[56] Y. Wang, S. Li, K. Smith, B.C. Waldman, A.S. Waldman, Intrachromosomal recombination between highly diverged DNA sequences is enabled in human cells deficient in Bloom helicase, DNA Repair 41 (2016) 73–84, 10.1016/j.dnarep.2016.03.005

[57] M.J. Wagner, J.A. Sharp, W.C. Summers, Nucleotide sequence of the thymidine kinase of herpes simplex virus type 1, Proc. Natl. Acad. Sci. U.S.A. 78 (1981) 1441–1445.

[58] K.M. Chapman, M.M. Wilkey, K.E. Potter, B.C. Waldman, A.S. Waldman, High homology is not required at the site of strand invasion during recombinational double-strand break repair in mammalian chromosomes, DNA Repair 60 (2017) 1–8, 10.1016/j.dnarep.2017.10.006

[59] B.C. Capel, F.S. Collins, Human laminopathies: nuclei gone genetically awry. Nat. Rev. Genet.7(2006):940-952, 10.1038/nrg1906

[60] E. Petermann M.L. Orta, N. Issaeva, N. Schultz, and T. Helleday, Hydroxyurea-stalled replication forks become progressively inactivated and require two different RAD51-mediated pathways for restart and repair, Mol. Cell. 37(2010) 492–502. 10.1016/j.molcel.2010.01.021.

[61] P. Scaffidi, and T. Misteli, Lamin A-dependent misregulation of adult stem cells associated with accelerated ageing, Nature Cell Biology 10 (2008) 452–459, 10.1038/ncb1708

[62] A. Chojnowski, P.F. Ong, M.X.R. Foo D. Liebl, L.-P. Hor, C.L. Stewart, and O. Oliver Dreesen, Heterochromatin loss as a determinant of progerin-induced DNA damage in Hutchinson–Gilford Progeria Aging Cell. 19 (2020) e13108, 10.1111/acel.13108

[63] A. Kychygina, M. Dall’Osto, J.A.M. Allen, J.-C. Cadoret, V. Piras, H.A. Picket, and L. Crabbe, Progerin impairs 3D genome organization and induces fragile telomeres by limiting the dNTP pools, Scientific Reports 11 (2021) 13195, 10.1038/s41598-021-92631-z

[64] . C.V. Wallis, A.N. Sheerin, M.H. Green, C.J. Jones, D. Kiplingand R.G. Faragher, Fibroblast clones from patients with Hutchinson–Gilford progeria can senesce despite the presence of telomerase, Exp. Gerontol.(2004) 39, 461–467.

[65] J. Mateos, J. Fafián-Labora, M. Morente-López, I. Lesende-Rodriguez, L. Monserrat, M.A. Ódena, E. Oliveira, J. de Toro, and M.C. Arufe, Next-generation sequencing and quantitative proteomics of Hutchinson-Gilford progeria syndrome-derived cells point to a role of nucleotide metabolism in premature aging, PLoS ONE 13 (2018) e0205878, 10.1371/journal.pone.0205878

[66] A. J. Lopez-Contreras, J. Specks, J.H. Barlow, C. Ambrogio, C. Desler, S. Vikingsson, S. Rodrigo-Perez, H. Green, L.J. Rasmussen, M. Murga, A. Nussenzweig, and O. Fernandez-Capetillo, Increased Rrm2 gene dosage reduces fragile site breakage and prolongs survival of ATR mutant mice, Genes and Dev. 29 (2015) 690–695, 10.1101/gad.256958.114

